# From ridge tops to ravines: landscape drivers of chimpanzee ranging patterns

**DOI:** 10.1101/795393

**Authors:** Samantha J. Green, Bryan J. Boruff, Cyril C. Grueter

## Abstract

Recent improvements in tracking technologies have resulted in a growing number of fine-scale animal movement studies in a variety of fields from wildlife management to animal cognition. Most studies assume that an animal’s “optimal” foraging route is linear, ignoring the role the energy landscape can play in influencing movement efficiency. Our objective was to investigate whether landscape features that affect movement costs; topographic variation, and super and substrate, influence the movement of chimpanzees (*Pan troglodytes*) in a rugged, montane environment. We tested for route re-use and preferential use of human-made trails and ridge tops using 14 months of focal follow data from 14 individuals and maps of established chimpanzee trails. Chimpanzees travelled on human-made trails significantly more than expected and showed weak preference for use of ridge tops for travel. Line density analysis demonstrated route re-use in chimpanzees and uncovered a network of high-use routes across their range. To our knowledge, this is the first study to empirically demonstrate route re-use and preferential use of human-made trails for travel by chimpanzees. We discuss the energetic and cognitive benefits of trail use and the implications for chimpanzee sociality. By applying the latest GIS analytical techniques to fine-scale movement data, this study demonstrates the importance of incorporating landscape features in predictive animal movement models.

## 1 Introduction

According to optimal foraging theory, natural selection favours animals that move across their environment in a way that minimizes energy expenditure while maximising energy gain (MacArthur and Pianka, 1966). As such, animals with good spatial cognition are able to choose the most efficient path between resources (Fagan et al., 2013). With recent improvements in tracking technologies, the collection and analysis of animal movement data in spatial cognition studies is growing (Janson and Byrne, 2007, Fagan et al., 2013, Garber and Dolins, 2014). However, few studies consider how landscape driven variation in movement costs, termed the energy landscape (Shepard et al., 2013, Wilson et al., 2012a), influences the choices animals make when moving about their environment (Leblond et al., 2010, Shepard et al., 2013, Howard et al., 2015, Strandburg-Peshkin et al., 2017). For example, linearity is a commonly used measure for travel efficiency (Valero and Byrne, 2007, Cunningham and Janson, 2007, Normand and Boesch, 2009, Asensio et al., 2011, Janmaat et al., 2013); however, this approach does not take into account landscape factors, such as topographic variability, which may result in more energy efficient sinuous routes. Therefore, identifying the role energy landscape plays in influencing animal movement patterns and path preferences is crucial to improving optimisation models of animal movement and thus our understanding of animal spatial cognition.

Shepard et al. (2013) identify three landscape factors that can influence energetic costs of terrestrial animal movement: topographic variation, and super and substrate penetrability. Large bodied animals exert significantly more energy travelling up slopes than on level ground (Halsey and White, 2017). The ratio of oxygen consumption to speed for a chimpanzee running up a 15 degree incline is double that of a chimpanzee running on level ground (Taylor et al., 1972). When drivers that favour increased speed over cost (e.g. predator avoidance and access to ephemeral food resources) are absent, animals living in rugged, steeply sloping, environments may therefore alter their ranging behaviour to avoid sharp inclines (Ganskopp et al., 2000, Dickson et al., 2005, Wall et al., 2006) and preferentially use ridges for travel due to their relatively straight curvature and shallow slopes (Surface-Evans and White, 2012, Pingel, 2010). Human hiking trails often follow ridges in mountainous areas (Pingel, 2010) and for example, indigenous people in the Amazon Basin prefer to hunt along ridge tops, as they view traversing valleys as too energetically costly (Yost and Kelley, 1983, Milton, 2000).

Substrate is defined as “the medium over or on which an animal moves” (Shepard et al., 2013 p. 299), and superstrate as “any material against which an animal must push to move” (Shepard et al., 2013 p. 300). Humans expend 1.7 to 2.7 times more energy walking on soft substrates such as sand than firmer substrates such as concrete (Pinnington and Dawson, 2001). For terrestrial animals, superstrate includes any material that extends above the substrate, with taller, denser and more rigid superstrates being more costly (Shepard et al., 2013). Studies have found that dense brush and snow are more energetically costly for humans to traverse than open ground (Soule and Goldman, 1975, Pandolf et al., 1976, Richmond et al., 2019), but there is little research on this topic for non-human mammals.

It can be difficult to study the movement costs (i.e. energy expenditure) associated with individual landscape factors as they often occur in combination. For example, human-made trails, paths and roads have compact level substrate, cleared superstrate, and are often designed to avoid steep slopes. Although most mammal species avoid human-made infrastructure (review in Benítez-López et al., 2010), researchers have documented preferential use of roads for travel by several North American mammals (cougars: Dickson et al. 2005; wolves: Thurber et al. 1994, Whittington et al., 2005; and grizzly bears: Roever et al., 2010). The energy savings offered by roads and human-made trails may outweigh the costs of exposure to humans for animals living in heterogeneous anthropogenic environments.

Terrestrial animals can also reduce travel costs by re-using their own tracks. This is a self-reinforcing behaviour with repeated route use resulting in greater substrate compaction and superstrate reduction. Therefore, the most frequently used routes offer the greatest savings in movement energy. Animals living in more variable environments may re-use routes of least-resistance at a high enough rate that they form well defined path-ways (Shepard et al., 2013, Perna and Latty, 2014).

There is growing evidence that naturally ranging chimpanzees (*Pan troglodytes*) possess good spatial knowledge of their environment (Normand et al., 2009, Normand and Boesch, 2009, Janmaat et al., 2013, Janmaat et al., 2014, Ban et al., 2016), making them an ideal species to examine landscape influences on ranging. Whilst landscape influences on fine-scale movement has to our knowledge not been documented in chimpanzees, the majority of chimpanzee research to date has been undertaken in relatively homogeneous landscapes which may not necessitate energy minimisation strategies.

Chimpanzee habitats occur over a wide altitudinal range, from tropical forests at sea-level in West Africa to high-altitude, montane forests in the Albertine Rift (Inskipp, 2005). Nyungwe National Park, a rugged montane forest in south-west Rwanda, supports a community of chimpanzees that range from 1 795 to 2 950 m above sea level (ASL), the highest known altitudinal limit of their species distribution. Nyungwe has relatively steep slopes and an open canopy that has created dense ground cover across much of the forest. The study community’s home range contains one major road, two smaller unpaved roads and a network of human-made trails (Figure 1). There is little research on the impact of super and substrate on the movement costs of non-human mammals, but we expect dense superstrates such as vine thickets and ground herbs will impede chimpanzee locomotion in Nyungwe and chimpanzees may minimise their daily energy expenditure by travelling on human-made trails or reusing their own tracks where possible. We also expect that the rugged terrain may drive preferential use of ridge tops for travel, as has been shown in several arboreal primates (Di Fiore and Suarez, 2007, Gregory et al., 2014, Presotto et al., 2018). To investigate whether landscape factors influence chimpanzee movements in this heterogenous environment, we test for: 1) repeated use of chimpanzee travel routes, 2) preferential use of human-made trails and roads for travel, and 3) preferential use of ridges for travel.

**Figure 1.**
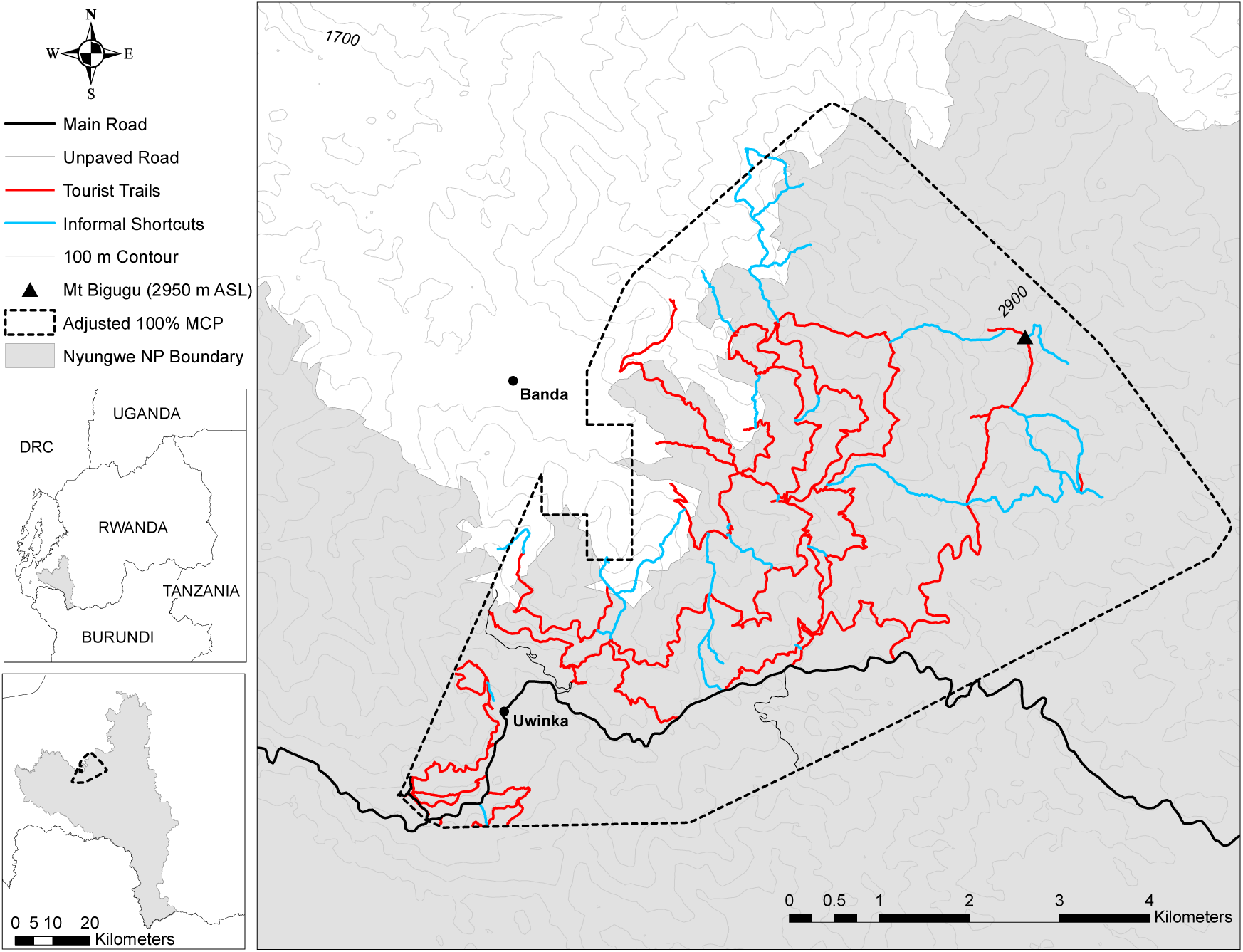
Location of the study area in Nyungwe National Park, Rwanda.

## 2 Methods

### 2.1 Study site and subjects

Nyungwe National Park protects 1 020 km^2^ of tropical montane forest in south-western Rwanda. Nyungwe receives 1 744 mm of annual rainfall with an average maximum and minimum temperature of 19.6°C and 10.9°C, respectively (Sun et al., 1996). The region experiences a major wet season from February to May, a long dry season from June to August, with a minor wet season from September to November and shorter dry season from December to January. The park is characterised by steep slopes with its highest peak measuring 2 950 m ASL (Fashing et al., 2007). The vegetation consists of both primary and secondary montane forest with a relatively open canopy that has resulted in a varied vegetation structure, with a dense herb layer in most areas (Fimbel et al., 2001, Plumptre et al., 2002).

Nyungwe is estimated to contain 380 individuals, the largest population of chimpanzees in Rwanda (IUCN, 2010). The Mayebe chimpanzee community have been habituated since 1997 by the Rwandan Development Board (formerly ORTPN) and Wildlife Conservation Society and are the focus of this study.

Data were collected between November 2016 and December 2017. By the end of our study, the Mayebe community consisted of 67 individuals: 14 adult and 4 sub-adult males, 18 adult and 7 sub-adult females, 12 juveniles and 12 infants (Smith and Green, 2018). We followed 14 individual adult male subjects over the course of this study.

### 2.2 Movement Observations

Sex differences in ranging patterns are common amongst eastern chimpanzees (*Pan troglodytes schweinfurthii*), with males traveling longer daily distances and having larger home ranges than anestrous females (Wrangham and Smuts, 1980, Chapman and Wrangham, 1993, Doran, 1997, Williams et al., 2002, Bates and Byrne, 2009). As such, to maximise route data collection only male chimpanzees were sampled. Behavioural data were collected on approximately ten days per month from November 2016 to December 2017. A focal adult male was randomly chosen from the first party encountered and followed for as long as possible, ideally until he built his night nest (focal animal sampling: Altman, 1974). Focal chimpanzee locations were recorded at 5 m intervals using a hand-held Garmin GPSMAP 64 device, with Global Navigation Satellite System (GLONASS) receiver. The Global Positioning System (GPS) accuracy was within 3–6 m throughout most of the Mayebe home range, but could increase to 20 m in some valleys due to poor signal. Focal party size and composition was recorded at 15 minute epochs. Individuals within 50 meters of each other were considered to be in the same party (following Clark and Wrangham, 1994).

We obtained 106 focal follows, with a mean duration of 8 h 38 min ± 2 h 23 min.

### 2.3 Analysis

#### 2.3.1 Home range

Several techniques for estimating home range size exist (Laver and Kelly, 2008). Minimum convex polygon (MCP, Hayne, 1949) and grid cell methods (Siniff and Tester, 1965, Haugen, 1942) are the most commonly used, but fixed kernel density estimation is considered by some authors to be the most accurate (Worton, 1987, Seaman and Powell, 1996) and local convex hull (LoCoH) is more capable of identifying hard boundaries, such as rivers, cliff edges or human settlements, that may not be used by the animal (Getz and Wilmers, 2004, Getz et al., 2007). We calculated home range using all four methods to allow comparison among sites as recommended by Boyle (in press). The MCP technique creates a bounding polygon around a set of data points using convex angles (Hayne, 1949). The grid cell method involves the identification of cells that contain at least one locational data point (Siniff and Tester, 1965, Haugen, 1942). We used 500 × 500 m grid cells to align with previous chimpanzee studies (Herbinger et al., 2001, Lehmann and Boesch, 2005, Amsler, 2009). The fixed kernel method calculates the density of location points within a defined neighbourhood. We calculated 99%, 95%, 75% and 50% fixed kernel estimates using the least squares cross validation (LSCV) method (Worton, 1989, Gitzen and Millspaugh, 2003) to define the bandwidth as it fit the data more closely than the reference (Worton, 1995) and plug-in estimator methods (Sheather and Jones, 1991) and was similar in fit to the bias cross validation method (Worton, 1989). To minimise autocorrelation between data points we used a subsample of four points per day (selecting the points taken closest to 6 am, 9 am, 12 pm and 3 pm, n = 551). The LoCoH is a nonparametric method that creates kernels using either a fixed number of points (*k*), or points with a fixed (*r*) or adaptive (*a*) sphere-of-influence (Getz et al., 2007). We used the *a*-LoCoH method as the estimated home range contained less “holes” than the *k-* and *r-*LoCoH methods (Getz et al., 2007). We used an *a* of 10 000 m as it was the maximum distance between any two location points. Home range estimates were generated using the Geospatial Modelling Environment ArcGIS extension (GME, Beyer, 2014) and the adehabitat statistical package (Clement, 2006) in R version 3.6.0 (R Core Team, 2019). As the Mayebe chimpanzees range near the boundary of the National Park, all home range estimates, except the *a-*LoCoH, included areas of unsuitable habitat within the neighbouring village. We used the adjusted polygon method (Grueter et al., 2009) to erase grid cells that were known to contain village areas from the home range polygons.

#### 2.3.2 Travel routes

Daily travel routes (n = 106) were created by joining each consecutive waypoint (mean waypoints collected per day = 425) with a straight-line segment in ArcMap 10.6.

#### 2.3.3 Mapping trails, roads and ridges

We used the ArcGIS Hydrology Toolset to extract ridge lines from 30 m resolution Shuttle Radar Topography Mission (SRTM) elevation data. We visually inspected the extracted ridge lines using Google Earth to identify any anomalies and mapped any ridge lines that had not been extracted following the technique described by Gregory et al. (2014).

To map human-made trails and roads we walked all trails and roads within the Mayebe chimpanzees home range, taking GPS readings every 5 m. This included both tourist trails and informal shortcuts, one paved and two unpaved roads. Established chimpanzee trails were mapped whenever chimpanzees were observed travelling on them.

#### 2.3.4 Landscape relationships

To examine whether chimpanzees showed repeated use of travel routes we used the ArcGIS Spatial Analyst Line Density tool. The Line Density tool calculates the density (km/km^2^) of travel routes in the defined neighbourhood of each output grid cell. Gregory et al. (2014) recommend adjusting the search radius to reflect the spread of the study group. Chimpanzees live in multifemale-multimale groups with fission-fusion dynamics in which temporary parties of variable size and composition unite and split throughout the day (Nishida, 1968, Aureli et al., 2008, Grueter et al., 2012). During the study period focal chimpanzees ranged alone or in parties of up to 22 individuals with distance’s up to 50 m between party members. To reflect these variable party sizes/spreads, the Line Density tool was run with search radii of 10-50 m in 10 m increments. Increasing the search radii from 10 m to 20 m identified additional repeatedly used routes, but further increases resulted in an increase in the width of existing routes only (Figure 2). The 50 m search radius is used in figures, as the wider travel routes are easier to visualise. We used all 14 months of data for this analysis (n = 106 daily travel routes) as the chimpanzees were observed ranging at opposite ends of their home range during the two months that were sampled twice, November and December.

**Figure 2.**
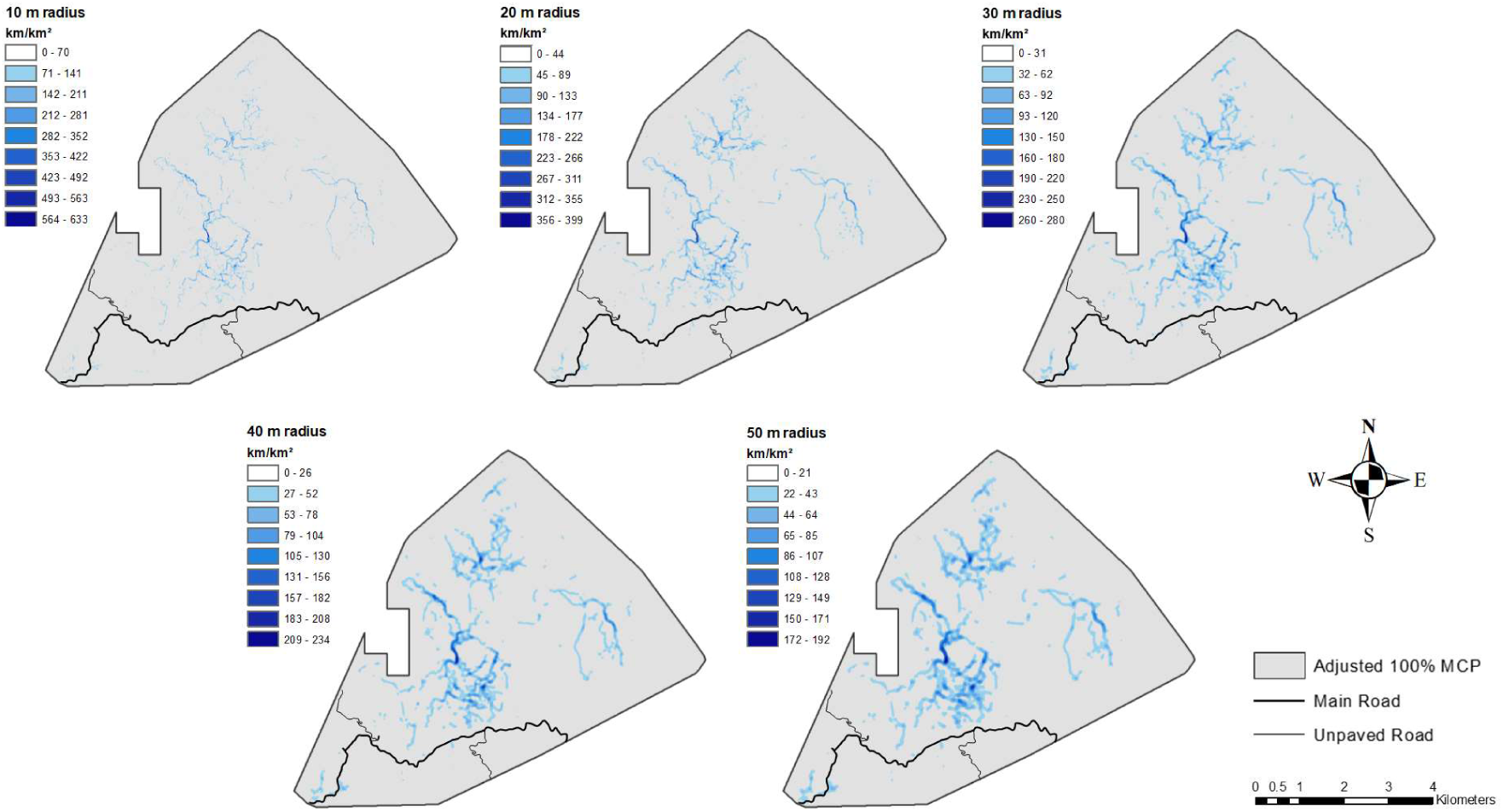
Chimpanzee travel routes density indicated by increasing search radii.

Preference for travel on human-made trails and ridges was analysed using third-order resource selection (Johnson, 1980). Landscape polygons were generated by buffering ridge lines by 50 m (to account for party spread) and trails by 10 m (as 10 m was the maximum width of trails). We then clipped the trail polygon to the ridge polygon to extract polygons that contained both trails and ridges. To estimate availability of landscape features, 5 000 random points were generated within the Mayebe home range (adjusted 100% MCP). The landscape category (trail, ridge, trail located on ridge or other) associated with each used (n = 4 083) locations from 14 individuals) and available location was queried in ArcMap. Landscape preference was analysed using a generalised linear mixed effects model with Binomial error structure. The dependent variable was 1 for used and 0 for available locations and the predictor variable was the landscape category (Boyce and McDonald, 1999, Manly et al., 2002). Available points were weighted by 5 000 prior to fitting the model (Fithian and Hastie, 2013). To account for certain individuals having a disproportionate effect on the dependent variable, the identity of the focal chimpanzee was included as a random effect (Gillies et al., 2006). The model coefficients (parameters, βi) are equivalent to selection ratios (Manly et al., 2002), with β_i_ > 0 indicating that chimpanzees use the landscape feature more than would be expected if used in proportion to its availability.

Model fitting was implemented in R version 3.6.0 (R Core Team, 2019) using the function ‘glmer’ of the package lme4 (version 1.1-21; Bates et al., 2015). Confidence intervals were computed using the function ‘confint’.

We did not test for the relationship between chimpanzee travel and roads statistically as chimpanzees were never observed travelling along roads, only crossing them.

## 3 Results

### 3.1 Home range

The Mayebe community home range size varied from 28.62 to 47.69 km^2^ depending on the method used (Table 1). As several daily routes extended beyond the 95% fixed kernel, and the grid cell and *a-*LoCoH methods excluded several areas that contained suitable habitat, we consider the adjusted 100% MCP estimate to be the most appropriate home range to use for our landscape analyses. The Mayebe chimpanzees ranged from 1 795 to 2 951 m ASL, the highest altitude recorded for chimpanzees.

**Table 1.**
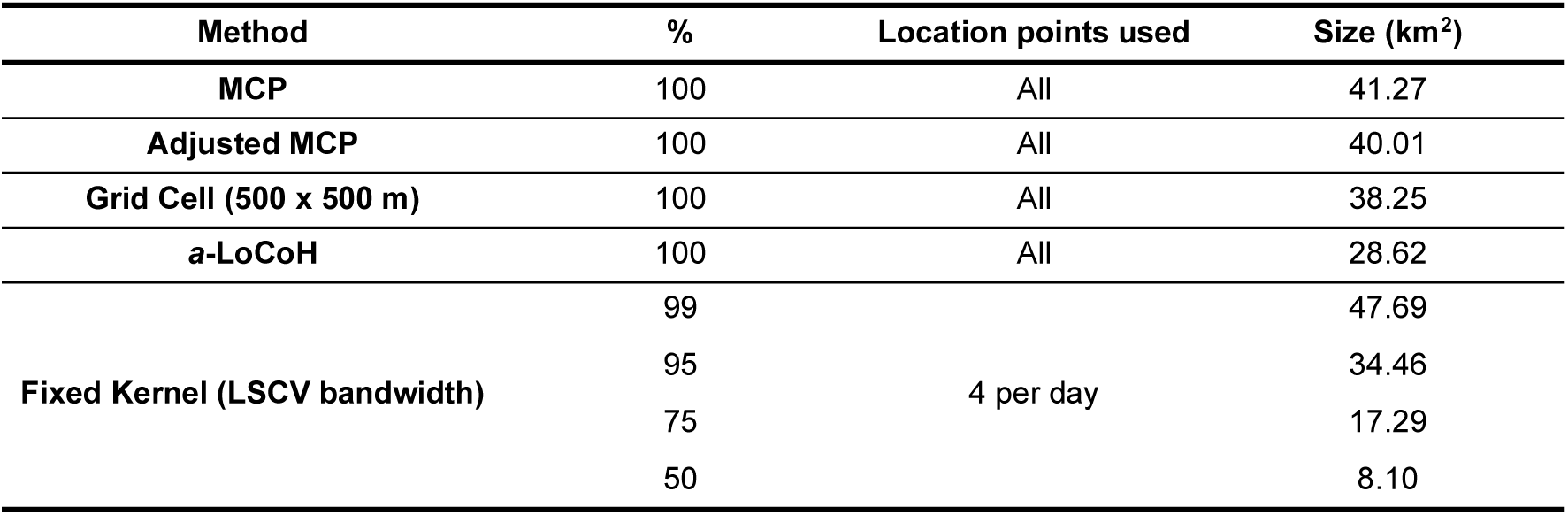
Home range estimates.

### 3.2 Landscape analysis

The route density plots demonstrate several repeatedly used routes across the Mayebe home range (Figure 2). The most intensely used routes occurred on human-made trails (Figure 3), but several were located outside of the trail network indicating that chimpanzees create their own trails. We recorded evidence of this in the field, with chimpanzees observed travelling along “established trails” that were not human-made on numerous occasions (Figure 3). These established trails were narrower than 1 metre, had level substrate that cut into steep slopes, were free from superstrate up to approximately 1 metre (sometimes forming a tunnel through vine thickets) and the bark had been worn off any dead logs or living vines that lay on the track, suggesting that these tracks were regularly used by chimpanzees. Established trails were relatively linear compared to human-made trails (Figure 3).

**Figure 3.**
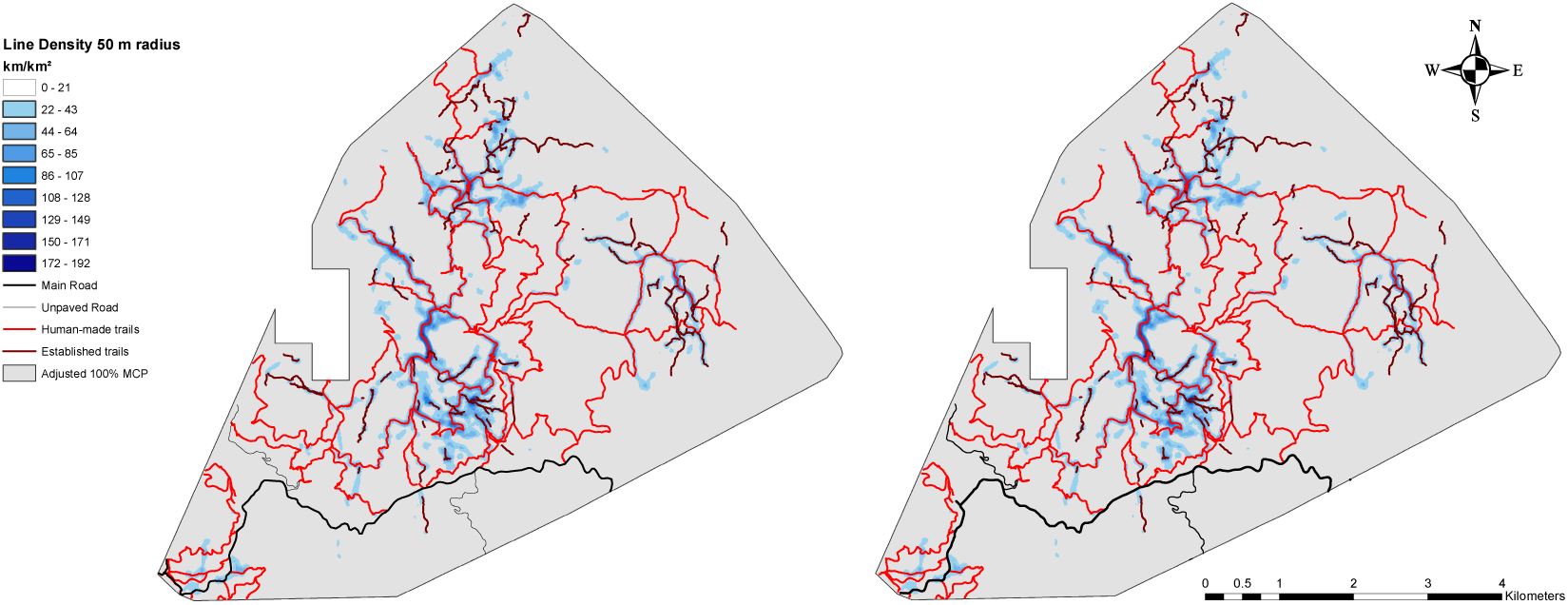
Chimpanzee travel route density showing overlap with human-made trails (left) and location of “established trails” (right). The 50 m search radius is used for visualisation purposes.

The most intensely used routes occurred on human-made trails (Figure 3) and chimpanzees did preferentially travel on human-made trails, whether they occurred on ridges (β_i_ = 2.75) or not (β_i_ = 2.78, Table 2), suggesting that chimpanzees preferentially use them for travel. Whilst some high-use routes and established trails fell on ridge tops, chimpanzees showed weak preference for travel on ridges (β_i_ =0.40, Table 2).

**Table 2.** Estimated selection coefficients for chimpanzee location points (n=45 083). SE: standard error; CI: confidence interval

| Variable | Estimate | SE | z | 95% CI |
| --- | --- | --- | --- | --- |
| Intercept | -8.69 | 0.076 |  |  |
| Ridge | 0.41 | 0.012 | 34.13 | 0.38, 0.43 |
| Trail | 2.78 | 0.013 | 215.14 | 2.75, 2.80 |
| Trails located on ridges | 2.75 | 0.014 | 201.77 | 2.72, 2.77 |

## 4 Discussion

Landscape features can have significant impacts on the movement efficiency of animals and thus directly influence their fitness (Wilson et al., 2012a, Perna and Latty, 2014). While large scale responses to energy landscape are well understood, there are few studies that consider finer scale responses and those that do generally only focus on a single factor, such as slope (Wilson et al., 2012a, Shepard et al., 2013). Here, we provide new insights into drivers of chimpanzee movement.

Our line density analysis uncovered a network of high-use chimpanzee movement routes across the Mayebe community’s home range. The most intensely used routes were located on human-made trails and statistical analysis showed that chimpanzees preferentially travel on human-made trails. Whilst chimpanzees have been observed using human-made trails in Budongo Forest Reserve, Uganda (Zommers et al., 2013), this study is the first to show chimpanzee *preference* for travel on human-made trails and adds to the few documented cases for non-human primates (e.g. baboons: Noser and Byrne, 2013; Strandburg-Peshkin et al., 2017). There are costs associated with travelling on human-made trails, including exposure to humans (Brncic et al., 2015) and the ‘turn costs’ associated with travelling on tortuous routes (Wilson et al., 2013, Halsey, 2016). Conversely, by travelling on human-made trails animals can decrease energy expenditure, due to the compact level substrate and cleared superstrate, without having to bear the cost of trail construction or maintenance (Perna and Latty, 2014). Several of Nyungwe’s trails also have bridges that facilitate travel across streams that would otherwise be difficult or impossible to traverse. The savings in movement energy and the chimpanzees’ habituation to humans are likely both important factors influencing their preferential use of human-made trails.

Chimpanzees showed weak preference for travel on ridge tops; however, several repeatedly used routes did follow ridgelines and we observed chimpanzees using ridges to travel up Nyungwe’s highest mountain on several occasions. The energetic benefits of travelling on ridges is likely influenced by the distance and elevation change between the start and end points of travel segments. For example, the cost of deviating to use a ridge may not outweigh the energetic gains when the difference in elevation starting point and goal is relatively small, but the deviation may be worthwhile for larger elevation changes. A more sophisticated model would be required to examine more nuanced influences.

While some high-use routes were associated with human-made trails and ridges, many were not associated with any prominent topographic features and some were orientated perpendicular to ridges, requiring travel up and down steep valleys. As there are no other large mammals in Nyungwe, we expect that these routes were likely formed by the same mechanisms that produce human pedestrian trails (Helbing et al., 1997a, Helbing et al., 1997b), whereby chimpanzees will preferentially use the existing human trail system, but when the detour is too large they will take the most efficient, or least-cost, route to their goal. In doing so they trample vegetation, compacting the substrate and reducing the superstrate, which makes the route more attractive to other chimpanzees. The more frequently the route is used the more attractive it becomes. When the frequency of use is greater than vegetation growth, the route is maintained and eventually becomes an established trail (Shepard et al., 2013, Perna and Latty, 2014).

Much of Nyungwe has been colonised by a rapidly growing climbing vine, *Sericostachys scandens* (Chao et al., 2009). However, chimpanzees were observed trampling through and subsequently reusing an 800 m route through *Sericostachys scandens* that was maintained for at least three weeks. In comparison to other primates that expand and contract their monthly ranges in response to changes in fruit availability (Hemingway and Bynum, 2005), the Mayebe community have a concentrated ranging pattern, with monthly home ranges that shift location across fruiting seasons, but remain relatively consistent in size (Moore et al., 2018, S.J. Green, unpublished data). This ranging strategy promotes more frequent trail re-use allowing for maintenance of seasonal trails.

Human trails in rugged environments are usually sinuous, as they are designed to avoid steep slopes. In comparison the established chimpanzee trails recorded in Nyungwe were relatively linear (Figure 3), seemingly ignoring the steep terrain. If established trails were routes that minimise metabolic costs (as suggested by Shepard et al., 2013), their relative linearity indicates that chimpanzees expend less energy travelling up slopes than humans. Chimpanzees will thus incur ‘turn costs’ (Wilson et al., 2013, Halsey, 2016) when travelling on human-made trails. The existence of relatively linear, high-use established trails and the preferential use of human-made trails despite ‘turn costs’ may indicate that substrate and superstrate are more important drivers of chimpanzee movement than slope, but more detailed elevation, super and substrate information are needed to test this. With fine-scale remote sensing technologies becoming increasingly more affordable, future studies could obtain this information using a combination of imagery collected from satellite and/or Unmanned Aerial Vehicles (UAV)’s and field surveys (see Strandburg-Peshkin et al., 2017 for an example). The relative importance of each landscape factor could then be tested using step-selection models (Thurfjell et al., 2014).

Other drivers of animal movement, such as predator avoidance (Boesch, 1991, Hodges et al., 2014), competition with neighbouring groups (Wilson et al., 2012b, Crofoot, 2013), mating opportunities (Clutton-Brock, 1989, Eberle and Kappeler, 2002) and within-group social interactions (Strandburg-Peshkin et al., 2017) may also be at play. For example, human-made and established trails appeared to facilitate some chimpanzee fusion events in Nyungwe. We regularly observed chimpanzees pant-hoot (a long-distance vocalisation) upon reaching a trail, at which point they would stop to wait for another foraging party to arrive before moving off together. The use of trails as “reunion sites” is consistent with Di Fiore and Suarez’s (2007) observations of spider monkey fusion events that commonly occur at key nodes in their arboreal route network. However, as dense undergrowth made it difficult to observe “off-trail” chimpanzee fusion events, it is not known whether chimpanzee fusion events were more common on trails than off. Reusing routes can also play an important role in maintaining contact between foraging parties. (Di Fiore and Suarez, 2007, Perna and Latty, 2014, Strandburg-Peshkin et al., 2017). Chimpanzees were often observed following an established trail that another party had taken several hours before. Regular use of human-made and established trails may thus help to maintain group cohesion in primates with flexible grouping patterns.

Without knowing the functional relationship between movement costs and terrain, it is difficult to isolate the energy landscape from other potential drivers of animal movement (Shepard et al., 2013, Lempidakis et al., 2018). However, insight can be gained by comparing the movement patterns of chimpanzees in different environments. Chimpanzee’s in Taï, Côte d’Ivoire rarely re-use routes (Jang et al., 2019). Taï has relatively flat terrain and sparse undergrowth (Boesch and Boesch-Achermann, 2000), which suggests that landscape factors are an important driver of chimpanzee route re-use. Studies of route re-use and preferential use of human-made trails by chimpanzee in habitats with dense undergrowth, such as Mahale, Tanzania (Nakamura et al., 2015) and Kahuzi-Biega, Democratic Republic of Congo (Yamagiwa and Basabose, 2006) are needed to shed light on the importance of superstrate in driving preferential trail use.

Our study also provides new insights into the spatial cognition of chimpanzees. The form of cognitive maps used by primates is a growing area of research that remains subject to debate (Garber and Dolins, 2014, Trapanese et al., 2018). Route re-use is often interpreted as evidence for use of a topological mental map (Milton, 2000, Di Fiore and Suarez, 2007, Noser and Byrne, 2007, Presotto and Izar, 2010, Hopkins, 2016). Normand and Boesch (2009) found evidence to support use of a Euclidean mental map by chimpanzees in Taï National Park, Côte d’Ivoire, but recommended further research in different habitat types. Euclidean maps may be necessary in relatively homogenous landscapes, such as Taï (Normand and Boesch, 2009), but not in more heterogenous habitats, such as Nyungwe, where the greater availability of landmarks make a topological representation of space more viable (Fagan et al., 2013). However, it is impossible to know for certain which representation of space Nyungwe chimpanzees use, as they gain the benefit of reduced energetic expenditure, as well as possible reductions in the cognitive demands of navigation (Dukas, 1999) by travelling on trails. It is possible that they use a mixture of both topological and Euclidean systems of reference, whereby they update their position with the aid of the configuration of trails and other landmarks (Benhamou, 1997).

Lastly, we limited our analysis to male chimpanzees as males were expected to use human-made trails and ridges more often, as they are usually more terrestrial (Wrangham and Smuts, 1980, Hunt, 1989, Doran, 1993, Takemoto, 2004, Pokempner, 2009), and travel longer distances per day (Wrangham and Smuts, 1980, Chapman and Wrangham, 1993, Doran, 1997, Williams et al., 2002, Bates and Byrne, 2009) and between food patches (Pokempner, 2009) than anestrous females. However, the energetic demands of gestation, lactation and travel with dependent offspring may make anestrous females more sensitive to the energy landscape than males. Female eastern chimpanzees also concentrate their ranging in small, individual core areas (Murray et al., 2007), which may promote more frequent route re-use. Future research on anestrous female ranging responses to the energy landscape may yield further insights into sex differences in foraging strategies.

### 4.1 Conclusion

Despite predictive models of animal movement being used in a growing number of studies with application to a variety fields from cognition to conservation planning (Kays et al., 2015), the influence of energy landscape on animal ranging decisions has been largely ignored. By utilising advances in GIS analytical techniques and mapping established trails, we demonstrate how landscape features can shape animal ranging patterns. To our knowledge, this is the first study to empirically demonstrate route re-use and preferential use of human-made trails by chimpanzees. These ranging strategies may be driven by the need to conserve energy whilst moving through dense superstrate in a rugged low resource montane environment. Further studies in varied environments would shed light on this.

## References

Altmann, J. (1974) ‘Observational study of behavior: sampling methods’, Behaviour, 49(3), pp. 227–266.

Amsler, S. J. (2009) Ranging behavior and territoriality in chimpanzees at Ngogo, Kibale National Park, Uganda. PhD Thesis, University of Michigan, Ann Arbor, Michigan.

Asensio, N., Brockelman, W. Y., Malaivijitnond, S. and Reichard, U. H. (2011) ‘Gibbon travel paths are goal oriented’, Animal Cognition, 14(3), pp. 395–405.

Aureli, F., Schaffner, C. M., Boesch, C., Bearder, S. K., Call, J., Chapman, C. A., Connor, R., Fiore, A. D., Dunbar, R. I. and Henzi, S. P. (2008) ‘Fission-fusion dynamics: new research frameworks’, Current Anthropology, 49(4), pp. 627–654.

Ban, S. D., Boesch, C., N’Guessan, A., N’Goran, E. K., Tako, A. and Janmaat, K. R. (2016) ‘Taï chimpanzees change their travel direction for rare feeding trees providing fatty fruits’, Animal Behaviour, 118, pp. 135–147.

Bates, D., Mächler, M., Bolker, B. and Walker, S. (2015) ‘Fitting linear mixed-effects models using lme4’, Journal of Statistical Software, 67(1), pp. 1–48.

Bates, L. A. and Byrne, R. W. (2009) ‘Sex differences in the movement patterns of free-ranging chimpanzees (Pan troglodytes schweinfurthii): foraging and border checking’, Behavioral Ecology and Sociobiology, 64(2), pp. 247–255.

Benhamou, S. (1997) ‘On systems of reference involved in spatial memory’, Behavioural Processes, 40(2), pp. 149–163.

Benítez-López, A., Alkemade, R. and Verweij, P. A. (2010) ‘The impacts of roads and other infrastructure on mammal and bird populations: A meta-analysis’, Biological Conservation, 143(6), pp. 1307–1316.

Beyer, H. L. (2014) Geospatial modelling environment for ArcGIS. Available at: http://www.spatialecology.com/gme.

Boesch, C. (1991) ‘The effects of leopard predation on grouping patterns in forest chimpanzees’, Behaviour, 117(3), pp. 220–241.

Boesch, C. and Boesch-Achermann, H. (2000) The chimpanzees of the Taï Forest: Behavioural ecology and evolution. New York: Oxford University Press.

Boyce, M. S. and McDonald, L. L. (1999) ‘Relating populations to habitats using resource selection functions’, Trends in Ecology & Evolution, 14(7), pp. 268–272.

Boyle, S. A. (in press) ‘Home range analysis: Why the methods matter’, in Shaffer, C.A., Dolins, F., Porter, L.L., Hickey, J.R. & Nibbelink., N.P. (eds.) GPS and GIS for Primatologists. Cambridge: Cambridge University Press.

Brncic, T., Amarasekaran, B., McKenna, A., Mundry, R. and Kühl, H. S. (2015) ‘Large mammal diversity and their conservation in the human-dominated land-use mosaic of Sierra Leone’, Biodiversity and conservation, 24(10), pp. 2417–2438.

Chao, N., Rugyerinyange, L. and Scholte, P. (2009) International Conference on the Impact of Sericostachys scandens on the Conservation of Nyungwe National Park (Rwanda): Protected Areas Biodiversity Project (PAB) Available at: https://rwanda.wcs.org/About-Us/Publications.aspx.

Chapman, C. A. and Wrangham, R. W. (1993) ‘Range use of the forest chimpanzees of Kibale: implications for the understanding of chimpanzee social organization’, American Journal of Primatology, 31(4), pp. 263–273.

Clark and Wrangham, R. W. (1994) ‘Chimpanzee arrival pant-hoots: Do they signify food or status?’, International Journal of Primatology, 15(2), pp. 185–205.

Clement, C. (2006) ‘The package “adehabitat” for the R software: a tool for the analysis of space and habitat use by animals’, Ecological Modelling, 197, pp. 516–519.

Clutton-Brock, T. H. (1989) ‘Mammalian mating systems’, Proceedings of the Royal Society of London. Series B. Biological sciences, 236(1285), pp. 339–372.

Crofoot, M. C. (2013) ‘The cost of defeat: Capuchin groups travel further, faster and later after losing conflicts with neighbors’, American journal of physical anthropology, 152(1), pp. 79–85.

Cunningham, E. and Janson, C. (2007) ‘Integrating information about location and value of resources by white-faced saki monkeys (*Pithecia pithecia*)’, Animal cognition, 10(3), pp. 293–304.

Di Fiore, A. and Suarez, S. (2007) ‘Route-based travel and shared routes in sympatric spider and woolly monkeys: cognitive and evolutionary implications’, Animal Cognition, 10(3), pp. 317–329.

Dickson, B. G., Jenness, J. S. and Beier, P. (2005) ‘Influence of vegetation, topography, and roads on cougar movement in Southern California’, Journal of Wildlife Management, 69(1), pp. 264–276.

Doran, D. (1993) ‘Comparative locomotor behavior of chimpanzees and bonobos: the influence of morphology on locomotion’, American Journal of Physical Anthropology, 91(1), pp. 83–98.

Doran, D. (1997) ‘Influence of seasonality on activity patterns, feeding behavior, ranging, and grouping patterns in Tai chimpanzees’, International Journal of Primatology, 18(2), pp. 183–206.

Dukas, R. (1999) ‘Costs of memory: ideas and predictions’, Journal of Theoretical Biology, 197(1), pp. 41–50.

Eberle, M. and Kappeler, P. M. (2002) ‘Mouse lemurs in space and time: a test of the socioecological model’, Behavioral Ecology and Sociobiology, 51(2), pp. 131–139.

Fagan, W. F., Lewis, M. A., Auger-Methe, M., Avgar, T., Benhamou, S., Breed, G., LaDage, L., Schlagel, U. E., Tang, W. W., Papastamatiou, Y. P., Forester, J. and Mueller, T. (2013) ‘Spatial memory and animal movement’, Ecol Lett, 16(10), pp. 1316–29.

Fashing, P. J., Mulindahabi, F., Gakima, J.-B., Masozera, M., Mununura, I., Plumptre, A. J. and Nguyen, N. (2007) ‘Activity and ranging patterns of Colobus angolensis ruwenzorii in Nyungwe Forest, Rwanda: possible costs of large group size’, International Journal of Primatology, 28(3), pp. 529–550.

Fimbel, C., Vedder, A., Dierenfeld, E. and Mulindahabi, F. (2001) ‘An ecological basis for large group size in Colobus angolensis in the Nyungwe Forest, Rwanda’, African Journal of Ecology, 39(1), pp. 83–92.

Fithian, W. and Hastie, T. (2013) ‘Finite-sample equivalence in statistical models for presence-only data’, The annals of applied statistics, 7(4), pp. 1917.

Ganskopp, D., Cruz, R. and Johnson, D. (2000) ‘Least-effort pathways: a GIS analysis of livestock trails in rugged terrain’, Applied Animal Behaviour Science, 68(3), pp. 179–190.

Garber, P. A. and Dolins, F. L. (2014) ‘Primate spatial strategies and cognition: Introduction to this special issue’, American Journal of Primatology, 76(5), pp. 393–398.

Getz, W. M., Fortmann-Roe, S., Cross, P. C., Lyons, A. J., Ryan, S. J. and Wilmers, C. C. (2007) ‘LoCoH: nonparameteric kernel methods for constructing home ranges and utilization distributions’, PloS one, 2(2), pp. e207.

Getz, W. M. and Wilmers, C. C. (2004) ‘A local nearest-neighbor convex-hull construction of home ranges and utilization distributions’, Ecography, 27(4), pp. 489–505.

Gillies, C. S., Hebblewhite, M., Nielsen, S. E., Krawchuk, M. A., Aldridge, C. L., Frair, J. L., Saher, D. J., Stevens, C. E. and Jerde, C. L. (2006) ‘Application of random effects to the study of resource selection by animals’, Journal of Animal Ecology, 75(4), pp. 887–898.

Gitzen, R. A. and Millspaugh, J. J. (2003) ‘Comparison of least-squares cross-validation bandwidth options for kernel home-range estimation’, Wildlife Society Bulletin, pp. 823–831.

Gregory, T., Mullett, A. and Norconk, M. A. (2014) ‘Strategies for navigating large areas: a GIS spatial ecology analysis of the bearded saki monkey, Chiropotes sagulatus, in Suriname’, American Journal of Primatology, 76(6), pp. 586–95.

Grueter, C. C., Chapais, B. and Zinner, D. (2012) ‘Evolution of multilevel social systems in nonhuman primates and humans’, International Journal of Primatology, 33(5), pp. 1002–1037.

Grueter, C. C., Li, D., Ren, B. and Wei, F. (2009) ‘Choice of analytical method can have dramatic effects on primate home range estimates’, Primates, 50(1), pp. 81.

Halsey, L. G. (2016) ‘Terrestrial movement energetics: current knowledge and its application to the optimising animal’, Journal of Experimental Biology, 219(10), pp. 1424–1431.

Halsey, L. G. and White, C. R. (2017) ‘A different angle: comparative analyses of whole-animal transport costs when running uphill’, Journal of Experimental Biology, 220(2), pp. 161–166.

Haugen, A. O. (1942) ‘Home range of the cottontail rabbit’, Ecology, 23(3), pp. 354–367.

Hayne, D. W. (1949) ‘Calculation of size of home range’, Journal of mammalogy, 30(1), pp. 1–18.

Helbing, D., Keltsch, J. and Molnar, P. (1997a) ‘Modelling the evolution of human trail systems’, Nature, 388(6637), pp. 47.

Helbing, D., Schweitzer, F., Keltsch, J. and Molnár, P. (1997b) ‘Active walker model for the formation of human and animal trail systems’, Physical review E, 56(3), pp. 2527.

Hemingway, C. A. and Bynum, N. (2005) ‘The influence of seasonality on primate diet and ranging’, in Brockman, D.K. & Van Schaik, C.P. (eds.) Seasonality in primates: Studies of living and extinct human and non-human primates. UK: Cambridge University Press.

Herbinger, I., Boesch, C. and Rothe, H. (2001) ‘Territory characteristics among three neighboring chimpanzee communities in the Taï National Park, Côte d’Ivoire’, International Journal of Primatology, 22(2), pp. 143–167.

Hodges, K. E., Cunningham, J. A. and Mills, L. S. (2014) ‘Avoiding and escaping predators: movement tortuosity of snowshoe hares in risky habitats’, Ecoscience, 21(2), pp. 97–103.

Hopkins, M. (2016) ‘Mantled howler monkey spatial foraging decisions reflect spatial and temporal knowledge of resource distributions’, Animal Cognition, 19(2), pp. 387–403.

Howard, A. M., Nibbelink, N. P., Madden, M., Young, L. A., Bernardes, S. and Fragaszy, D. M. (2015) ‘Landscape influences on the natural and artificially manipulated movements of bearded capuchin monkeys’, Animal Behaviour, 106, pp. 59–70.

Hunt, K. (1989) Positional behavior in Pan troglodytes at the Mahale Mountains and the Gombe Stream National Parks, Tanzania. PhD Thesis, University of Michigan.

Inskipp, T. (2005) ‘Chimpanzees (*Pan troglodytes*)’, in Caldecott, J. & Miles, L. (eds.) World Atlas of Great Apes and Their Conservation. Berkley: University of California Press, pp. 53–81.

IUCN (2010) Eastern Chimpanzee Management Plan (Pan troglodytes schweinfurthii): Eastern Chimpanzee Management Plan.

Jang, H., Boesch, C., Mundry, R., Ban, S. D. and Janmaat, K. R. (2019) ‘Travel linearity and speed of human foragers and chimpanzees during their daily search for food in tropical rainforests’, Scientific reports, 9.

Janmaat, K. R. L., Ban, S. D. and Boesch, C. (2013) ‘Chimpanzees use long-term spatial memory to monitor large fruit trees and remember feeding experiences across seasons’, Animal Behaviour, 86(6), pp. 1183–1205.

Janmaat, K. R. L., Polansky, L., Ban, S. D. and Boesch, C. (2014) ‘Wild chimpanzees plan their breakfast time, type, and location’, Proceedings of the National Academy of Sciences, 111(46), pp. 16343–16348.

Janson, C. and Byrne, R. (2007) ‘What wild primates know about resources: opening up the black box’, Animal cognition, 10(3), pp. 357–367.

Johnson, D. H. (1980) ‘The comparison of usage and availability measurements for evaluating resource preference’, Ecology, 61(1), pp. 65–71.

Kays, R., Crofoot, M. C., Jetz, W. and Wikelski, M. (2015) ‘Terrestrial animal tracking as an eye on life and planet’, Science, 348(6240).

Laver, P. N. and Kelly, M. J. (2008) ‘A critical review of home range studies’, The Journal of Wildlife Management, 72(1), pp. 290–298.

Leblond, M., Dussault, C. and Ouellet, J.-P. (2010) ‘What drives fine-scale movements of large herbivores? A case study using moose’, Ecography, 33(6), pp. 1102–1112.

Lehmann, J. and Boesch, C. (2005) ‘Bisexually bonded ranging in chimpanzees (Pan troglodytes verus)’, Behavioral Ecology and Sociobiology, 57(6), pp. 525–535.

Lempidakis, E., Wilson, R. P., Luckman, A. and Metcalfe, R. S. (2018) ‘What can knowledge of the energy landscape tell us about animal movement trajectories and space use? A case study with humans’, Journal of theoretical biology, 457, pp. 101–111.

MacArthur, R. H. and Pianka, E. R. (1966) ‘On optimal use of a patchy environment’, Animal Nature, 100, pp. 603–609.

Manly, B. F. J., McDonald, L. L., Thomas, D. L., McDonald, T. L. and Erickson, W. P. (2002) Resource selection by animals: statistical analysis and design for field studies. Dordrecht, The Netherlands: Kluwer.

Milton, K. (2000) ‘Quo vadis? Tactics of food search and group movement in primates and other animals’, in Boinski, S. & Garber, P.A. (eds.) On the move: How and why animals travel in groups. Chicago: University of Chicago Press, pp. 375–417.

Moore, J. F., Mulindahabi, F., Gatorano, G., Niyigaba, P., Ndikubwimana, I., Cipolletta, C. and Masozera, M. K. (2018) ‘Shifting through the forest: home range, movement patterns, and diet of the eastern chimpanzee (Pan troglodytes schweinfurthii) in Nyungwe National Park, Rwanda’, American journal of primatology, 80(8), pp. e22897.

Murray, C. M., Mane, S. V. and Pusey, A. E. (2007) ‘Dominance rank influences female space use in wild chimpanzees, Pan troglodytes: towards an ideal despotic distribution’, Animal Behaviour, 74(6), pp. 1795–1804.

Nakamura, M., Hosaka, K., Itoh, N. and Zamma, K. (2015) Mahale chimpanzees: 50 years of research. Cambridge University Press.

Nishida, T. (1968) ‘The social group of wild chimpanzees in the Mahali Mountains’, Primates, 9(3), pp. 167–224.

Normand, E., Ban, S. D. and Boesch, C. (2009) ‘Forest chimpanzees (Pan troglodytes verus) remember the location of numerous fruit trees’, Animal Cognition, 12(6), pp. 797–807.

Normand, E. and Boesch, C. (2009) ‘Sophisticated Euclidean maps in forest chimpanzees’, Animal Behaviour, 77(5), pp. 1195–1201.

Noser, R. and Byrne, R. W. (2007) ‘Mental maps in chacma baboons (Papio ursinus): using inter-group encounters as a natural experiment’, Animal Cognition, 10(3), pp. 331–340.

Noser, R. and Byrne, R. W. (2013) ‘Change point analysis of travel routes reveals novel insights into foraging strategies and cognitive maps of wild baboons’, American Journal of Primatology, 76(5), pp. 399–409.

Pandolf, K. B., Haisman, M. F. and Goldman, R. F. (1976) ‘Metabolic energy expenditure and terrain coefficients for walking on snow’, Ergonomics, 19, pp. 683–90.

Perna, A. and Latty, T. (2014) ‘Animal transportation networks’, Journal of The Royal Society Interface, 11(100), pp. 20140334.

Pingel, T. J. (2010) ‘Modeling slope as a contributor to route selection in mountainous areas’, Cartography and Geographic Information Science, 37(2), pp. 137–148.

Pinnington, H. C. and Dawson, B. (2001) ‘The energy cost of running on grass compared to soft dry beach sand’, Journal of Science and Medicine in Sport, 4(4), pp. 416–430.

Plumptre, A. J., Masozera, M., Fashing, P. J., McNeilage, A., Ewango, C., Kaplin, B. and Liengola, I. (2002) Biodiversity Surveys of the Nyungwe Forest Reserve in S.W. Rwanda. WCS Working Papers No. 18, Bronx, NY: Wildlife Conservation Society.

Pokempner, A. A. (2009) Fission -fusion and foraging: Sex differences in the behavioral ecology of chimpanzees (Pan troglodytes schweinfurthii). PhD Thesis, Stony Brook University.

Presotto, A. and Izar, P. (2010) ‘Spatial reference of black capuchin monkeys in Brazilian Atlantic Forest: egocentric or allocentric?’, Animal Behaviour, 80(1), pp. 125–132.

Presotto, A., Verderane, M. P., Biondi, L., Mendonca-Furtado, O., Spagnoletti, N., Madden, M. and Izar, P. (2018) ‘Intersection as key locations for bearded capuchin monkeys (Sapajus libidinosus) traveling within a route network’, Animal Cognition, 21(3), pp. 393–405.

R Core Team (2019) R: A language and environment for statistical computing. Vienna, Austria: R Foundation for Statistical Computing.

Richmond, P. W., Potter, A. W., Looney, D. P. and Santee, W. R. (2019) ‘Terrain coefficients for predicting energy costs of walking over snow’, Applied ergonomics, 74, pp. 48–54.

Roever, C. L., Boyce, M. S. and Stenhouse, G. B. (2010) ‘Grizzly bear movements relative to roads: application of step selection functions’, Ecography, 33(6), pp. 1113–1122.

Seaman, D. E. and Powell, R. A. (1996) ‘An evaluation of the accuracy of kernel density estimators for home range analysis’, Ecology, 77(7), pp. 2075.

Sheather, S. J. and Jones, M. C. (1991) ‘A reliable data-based bandwidth selection method for kernel density estimation’, Journal of the Royal Statistical Society: Series B (Methodological), 53(3), pp. 683–690.

Shepard, E. L., Wilson, R. P., Rees, W. G., Grundy, E., Lambertucci, S. A. and Vosper, S. B. (2013) ‘Energy landscapes shape animal movement ecology’, Am Nat, 182(3), pp. 298–312.

Siniff, D. B. and Tester, J. R. (1965) ‘Computer analysis of animal-movement data obtained by telemetry’, Bioscience, 15(2), pp. 104–108.

Smith, B. and Green, S. (2018) Mayebe Chimpanzee Community Composition and Identification, Nyungwe Forest, Rwanda, University of Western Australia.

Soule, R. G. and Goldman, R. F. (1975) ‘Terrain coefficients for energy cost prediction’, Journal of Applied Physiology, 32(5), pp. 706–708.

Strandburg-Peshkin, A., Farine, D. R., Crofoot, M. C. and Couzin, I. D. (2017) ‘Habitat and social factors shape individual decisions and emergent group structure during baboon collective movement’, Elife, 6.

Sun, C., Kaplin, B. A., Kristensen, K. A., Munyaligoga, V., Mvukiyumwami, J., Kajondo, K. K. and Moermond, T. C. (1996) ‘Tree phenology in a tropical montane forest in Rwanda’, Biotropica, pp. 668–681.

Surface-Evans, S. L. and White, D. A. (2012) ‘An introduction to the least cost analysis of social landscapes’, Least cost analysis of social landscape: archaeological case studies. The University of Utah Press. Salt Lake City, pp. 1–7.

Takemoto, H. (2004) ‘Seasonal change in terrestriality of chimpanzees in relation to microclimate in the tropical forest’, American Journal of Physical Anthropology, 124(1), pp. 81–92.

Taylor, C. R., Caldwell, S. L. and Rowntree, V. J. (1972) ‘Running Up and Down Hills: Some Consequences of Size’, Science, 178(4065), pp. 1096–1097.

Thurber, J. M., Peterson, R. O., Drummer, T. D. and Thomasma, S. A. (1994) ‘Gray wolf response to refuge boundaries and roads in Alaska’, Wildlife Society Bulletin, 22(1), pp. 61.

Thurfjell, H., Ciuti, S. and Boyce, M. S. (2014) ‘Applications of step-selection functions in ecology and conservation’, Movement ecology, 2(1), pp. 4.

Trapanese, C., Meunier, H. and Masi, S. (2018) ‘What, where and when: spatial foraging decisions in primates’, Biological Reviews.

Valero, A. and Byrne, R. (2007) ‘Spider monkey ranging patterns in Mexican subtropical forest: do travel routes reflect planning?’, Animal Cognition, 10(3), pp. 305–315.

Wall, J., Douglas-Hamilton, I. and Vollrath, F. (2006) ‘Elephants avoid costly mountaineering’, Current Biology, 16(14), pp. R527–R529.

Whittington, J., Colleen Cassady St, C. and Mercer, G. (2005) ‘Spatial responses of wolves to roads and trails in mountain valleys’, Ecological Applications, 15(2), pp. 543–553.

Williams, J. M., Pusey, A. E., Carlis, J. V., Farm, B. P. and Goodall, J. (2002) ‘Female competition and male territorial behaviour influence female chimpanzees’ ranging patterns’, Animal Behaviour, 63(2), pp. 347–360.

Wilson, R. P., Quintana, F. and Hobson, V. J. (2012a) ‘Construction of energy landscapes can clarify the movement and distribution of foraging animals’, Proceedings of the Royal Society of London B: Biological Sciences, 279(1730), pp. 975–980.

Wilson, M. L., Kahlenberg, S. M., Wells, M. and Wrangham, R. W. (2012b) ‘Ecological and social factors affect the occurrence and outcomes of intergroup encounters in chimpanzees’, Animal Behaviour, 83(1), pp. 277–291.

Wilson, R., Griffiths, I., Legg, P., Friswell, M., Bidder, O., Halsey, L., Lambertucci, S. A. and Shepard, E. (2013) ‘Turn costs change the value of animal search paths’, Ecology Letters, 16(9), pp. 1145–1150.

Worton, B. J. (1987) ‘A review of models of home range for animal movement’, Ecological Modelling, 38(3-4), pp. 277–298.

Worton, B. J. (1989) ‘Kernel methods for estimating the utilization distribution in home-range studies’, Ecology, 70(1), pp. 164–168.

Worton, B. J. (1995) ‘Using Monte Carlo simulation to evaluate kernel-based home range estimators’, The Journal of wildlife management, pp. 794–800.

Wrangham, R. W. and Smuts, B. B. (1980) ‘Sex differences in the behavioural ecology of chimpanzees in the Gombe National Park, Tanzania’, Journal of Reproduction and Fertility, pp. 13–31.

Yamagiwa, J. and Basabose, A. K. (2006) ‘Effects of fruit scarcity on foraging strategies of sympatric gorillas and chimpanzees’, in Hohmann, G., Robbins, M.M. & Boesch, C. (eds.) Feeding Ecology in Apes and Other Primates. Cambridge: Cambridge University Press, pp. 73–96.

Yost, J. A. and Kelley, P. M. (1983) ‘Shotguns, Blowguns, and Spears: The Analysis of Technological Efficiency’, Adaptive Responses of Native Amazonians. New York: Academic Press, pp. 189–224.

Zommers, Z., Macdonald, D. W., Johnson, P. J. and Gillespie, T. R. (2013) ‘Impact of human activities on chimpanzee ground use and parasitism (*Pan troglodytes*)’, Conservation letters, 6(4), pp. 264–273.

